# SpecGlob: rapid and accurate alignment of mass spectra differing from their peptide models by several unknown modifications

**DOI:** 10.1101/2022.05.31.494131

**Authors:** Albane Lysiak, Guillaume Fertin, Géraldine Jean, Dominique Tessier

## Abstract

**Background:** In proteomics, mass spectra representing peptides carrying multiple unknown modifications are particularly difficult to interpret, which results in a large number of unidentified spectra.

**Methods:** We developed SpecGlob, a dynamic programming algorithm that aligns pairs of spectra, each such pair being a Peptide-Spectrum Match (PSM) provided by any Open Modification Search (OMS) method. For each PSM, SpecGlob computes the best alignment according to a given score system, while interpreting the mass delta within the PSM as one or several unspecified modification(s). All alignments are provided in a file, written in a specific syntax.

**Results:** Using several sets of simulated spectra generated from the human proteome, we demonstrate that running SpecGlob as a post-analysis of an OMS method can significantly increase the number of correctly interpreted spectra, as SpecGlob is able to correctly and rapidly align spectra that differ by one or more modification(s) without any *a priori*.

**Conclusion:** Since SpecGlob explores all possible alignments that may explain the mass delta within a PSM, it reduces interpretation errors generated by incorrect assumptions about the modifications present in the sample or the number and the specificities of modifications carried by peptides. Our results demonstrate that SpecGlob should be relevant to align experimental spectra, although this consists in a more challenging task.

## 1. Introduction

Most of the spectra generated by mass spectrometry proteomic experiments remain unidentified after their analysis by an identification software. One of the main reasons for this poor identification rate is that most of these spectra correspond to the fragmentation of peptides carrying modifications (Bogdanow *et al*. 2016; Griss *et al*. 2016). In conventional approaches, identification methods try to pair imperfect experimental spectra to ideal reference spectra – called *theoretical spectra* – generated from a peptide database (Noor *et al*. 2021).

However, in order to avoid excessive computations, these methods limit the mass difference Δ*m* between any experimental spectrum and the theoretical spectra that will be tentatively paired to it – usually, Δ*m* is limited to the mass spectrometer tolerance. As a result, many potential Peptide Spectrum Matches (PSMs) involving modifications cannot be detected, since these modifications may imply a larger Δ*m*.

In order to overcome these limitations, Open Modification Search (OMS) methods allow matches between similar spectra that represent distinct chemical compounds with unequal masses whose difference may be large. Pioneering approaches developed since the early 2000s rely on the alignment of paired spectra (Pevzner *et al*. 2000; Pevzner *et al*. 2001; Tsur *et al*. 2005; Cliquet *et al*. 2009) using a dynamic programming algorithm (Bellman 1952). Their main limitation is their excessive execution time, explained by the need to compare a very large number of theoretical spectra per experimental spectrum. Therefore, subsequent methods put forward a limitation on the number of PSMs to be evaluated, by pre-selecting a subset of theoretical spectra for each experimental spectrum. Selecting paired spectra that exhibit a Δ*m* corresponding to a known modification or a combination of known (or most frequent) modifications is one way to limit the number of PSMs (Savitski *et al*. 2006; Solntsev *et al*. 2018; Na *et al*. 2019). The main drawback, however, is that only known modifications can be identified and their number per spectrum is limited. Alternative methods select theoretical spectra that contain small amino acids tags deduced *de novo* from experimental spectra (Searle *et al*. 2005; Na *et al*. 2012; Chi *et al*. 2018; Devabhaktuni *et al*. 2019; Na *et al*. 2019).

More recently, the study of Chick *et al*. (Chick *et al*. 2015) reboosted interest in methods that select pairs of correlated spectra, highlighting a large variety of modifications present in samples, all in a reasonable execution time. In the wake of this study, several strategies were explored with success to drastically decrease the execution time. Some choose to compare experimental spectra to spectral libraries (Horlacher *et al*. 2016; Burke *et al*. 2017; Kong *et al*. 2017; Bittremieux *et al*. 2019) that are smaller than the entire proteome and supposedly more representative of the expressed proteins. Conversely, others are still based on a comparison with the complete theoretical proteome (David *et al*. 2017; Kong *et al*. 2017).

Although a recent study attributes some advantage, in terms of accuracy and sensibility, to OMS methods based on spectra comparisons, their interpretation of spectra when multiple modifications occur, is still challenging (Riffle *et al*. 2022). Detecting modifications within a PSM is equivalent to finding an alignment between the paired spectra, accompanied with one or several mass offset(s) allowing to better align the peaks of the theoretical spectrum to those of the experimental spectrum. When only *one modification* differentiates both spectra (i.e., a subset of peaks is shifted by Δ*m*), or when the possible modifications are known in advance, the alignment problem is already addressed by several efficient algorithms (see ((Chalkley and Clauser 2012) for a review and (Shteynberg *et al*. 2019; Cifani *et al*. 2021) for new approaches). However, when the two spectra of a PSM are separated by *several modifications*, Δ*m* must be split into several parts, a complex operation especially without any *a priori* on the nature of the modifications the experimental spectrum carries. Some methods evaluate the most frequent modifications in the sample, and try to interpret Δ*m* by a combination of these frequent modifications (An *et al*. 2018; Geiszler *et al*. 2021). Unfortunately, this strategy mainly interprets artefacts, which are the most frequent modifications without any enrichment technique applied on samples. Moreover, given that these methods combine known modifications, they can possibly miss important biological modifications. We consider that low abundance modifications are essential targets to OMS methods given their potential biological importance.

We have therefore developed a new method, called SpecGlob, that generates alignments of PSMs, even when *several unknown modifications* differentiate the paired spectra. SpecGlob relies on dynamic programming as in some of the mentioned prior works; however, its objective greatly differs. Indeed, its goal is not to obtain PSMs; instead, for each given PSM (pre-computed by an OMS method chosen by the user), SpecGlob detects, without any *a priori* on their nature, the modifications that explain the differences between the two spectra. For each PSM, SpecGlob returns its best alignment under the form of a string we called *StModified*, that gives several indications on how to align the peptide to the spectrum.

We then evaluate the behaviour of SpecGlob on several simulated spectra datasets of increasing complexity. Firstly, after targeting modifications on specific amino acids, we compare our results to those obtained with the state-of-the-art method MODPlus, the recent version of MODa (Na *et al*. 2012; Na *et al*. 2019). MODPlus selects candidate peptides that share sequence tags with *Se*, and next aligns the spectra with a dynamic programming approach using an input modification list (limited to the modification database Unimod (Creasy and Cottrell 2004) or to a user-defined list). Secondly, we extend the dictionary of modifications taken into account, possibly involving several amino acids. To this end, we pairwise compare all theoretical spectra generated from the human proteome while excluding self-identifications. This allows us to simulate, in the reference database, the systematic absence of the peptide at the origin of any simulated experimental spectrum. Then, as in (Lysiak *et al*. 2021), we also introduce a classification of all *StModified* sequences that evaluates the level of difficulty of exactly retrieving the original amino acids sequence of a simulated experimental spectrum starting from its *StModified* sequence. Based on this classification, we investigate the strengths and weaknesses of SpecGlob, and conclude that SpecGlob behaves very well on all tested datasets. Finally, we discuss its use in an experimental context.

## 2. Materials and Methods

### 2.1 Spectra datasets and protein database

#### Ethical Compliance

All procedures performed in this study were in accordance with the ethical standards of the institutional and/or national research committee and with the 1964 Helsinki Declaration and its later amendments or comparable ethical standards.

All spectra datasets were generated from the human proteome, downloaded from Ensembl99, release GrCh38 (Yates *et al*. 2020). While the generation of spectra in SpecOMS is independent from the role a peptide plays (experimental (*Se*) or theoretical (*St*)), this generation differs through SpecGlob. When a peptide plays the role of *St*, it is modeled by a spectrum only containing *b*-ions, which is enough to represent the amino acids (because *y*-ions are complementary to *b*-ions). When a peptide plays the role of *Se*, it is considered as coming from an unknown peptide. Consequently, as one cannot differentiate the *b*-ions from the *y*-ions before any identification, both types of peaks are generated. Firstly, we generated two simulated sets of 50,000 theoretical spectra to compare SpecGlob to MODPlus. Tryptic unique peptides whose lengths ∈ [12,25] are transformed into simulated spectra carrying modifications on some amino acids as follows: in dataset *DatasetND*, asparagines were deamidated (N+0.984016) and sodium adducts were added on each aspartic acid (D+21.981943); in *dataset DatasetSCT*, serines were substituted by alanines (S-15.99), cysteines were carbamidomethylated (C+57.0214) and threonines were deleted (T-101.0477).

Secondly, in dataset *AllvsAll*, all human tryptic peptides whose lengths ∈[7,30] were selected to play the role of experimental spectra. The protein database merges *target* and *decoy* protein sequences (generated by reversing sequences).

### 2.2 PSMs generation with SpecOMS (parameters setting in Supplementary Data, section 1)

SpecOMS (David *et al*. 2017) is a very fast OMS identification software used to identify experimental spectra by comparison with theoretical spectra, based on the number of shared peaks (Shared Peaks Count, or SPC) between pairs of spectra. Given *Se*, SpecOMS selects all *St* that share at least *p* peaks with *Se* (*p* set by the user). Next, in order to produce the best PSM (*Se, St*), the experimental spectrum that shares the maximum number of peaks is chosen according to the shifted SPC method– see (Lysiak *et al*. 2021) for details.

### 2.3 SpecGlob algorithm (pseudocode in Supplementary Data, section 2)

SpecGlob relies on a dynamic programming approach to find the best alignment between the *N* masses (corresponding to *b*-ions) of *St* and the *M* masses (corresponding to both *b*- and *y*-ions) of *Se*, which are respectively stored in arrays *StMasses*[] and *SeMasses*[].

Since PSMs are produced by an OMS method, pairs of spectra are expected to share some similarities. For each PSM, SpecGlob looks for an alignment that maximizes a certain (user-defined) score, possibly splitting the mass difference Δ*m* between *Se* and *St* into several mass offsets. This score globally takes (positively) into account the number of aligned amino acids and to a lesser extent, the number of inserted mass offsets, and (negatively) the number of non-aligned amino acids. It is important to note that the number of mass offsets required for an optimal alignment is not defined or limited in advance.

A matrix *D* of size *N* × *M* is filled according to a score system and a set of rules (Equation (1)) which allow to compute, for any 0 ≤ *i* ≤ *N* − 1 and any 0 ≤ *j* ≤ *M* − 1, the value of *D*[*i*][*j*].

**Equation 1:**

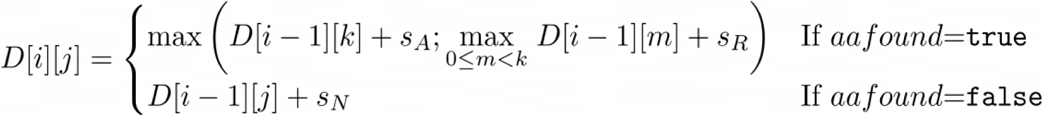

*D*[*i*][*j*] depends on a boolean, *aafound*, which is set to true only if the mass of the *i*-th amino acid of *St*, say *aa*, is found in *Se*. More precisely, *aafound*=true if there exists *k < j* such that the following holds:

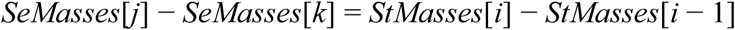

Otherwise, *aafound*=false.

The above formula uses three different score values, namely *s*_*A*_, *s*_*R*_ and *s*_*N*_ (where *A* stands for Align, *R* for Realign and *N* for No-align) to determine *D*[*i*][*j*] as follows (Equation (1)):

- if *aafound* is true, then:
  – if *aa* is found without a mass offset, the alignment comes from cell *D*[*i* − 1][*k*], and the corresponding score is *D*[*i* − 1][*k*] + *s*_*A*_;
  – if *aa* is found with a mass offset, the corresponding score is *D*[*i* − 1][*m*]+ *s*_*R*_, where *m* is the index (ranging from 0 to *k* − 1) that maximizes the score. If several such indices *m* exist, the largest is chosen.

The maximum value between the two above hypotheses is selected.

- if *aafound* is false, *D*[*i*][*j*] = *D*[*i* − 1][*j*] + *s*_*N*_.

In this study, *s*_*A*_ = 5, *s*_*R*_ = 2 and *s*_*N*_ = −4.

*D*[0][*j*] = 0 for every 0 ≤ *j* ≤ *M* −1 and for every 0 ≤ *i* ≤ *N* − 1, *D*[*i*][0] is set to *i* · *s*_*N*_, to mark amino acids as not found when the alignment starts.

All cells *D*[*i*][*j*] are then computed for *i* from 1 to *N* − 1 and for *j* from 1 to *M* − 1, i.e. from left to right and top to bottom, according to Equation (1). In the process, we store the origin (i.e. the coordinates) of the current cell’s score, together with the corresponding type of alignment (Align, Realign or No-Align), in a matrix called *Origin*.

Once *D* is filled, a backtrack algorithm is performed (Supplementary Data, section 2.4). An overview of the SpecGlob workflow is provided Figure 1.

**Figure 1:**
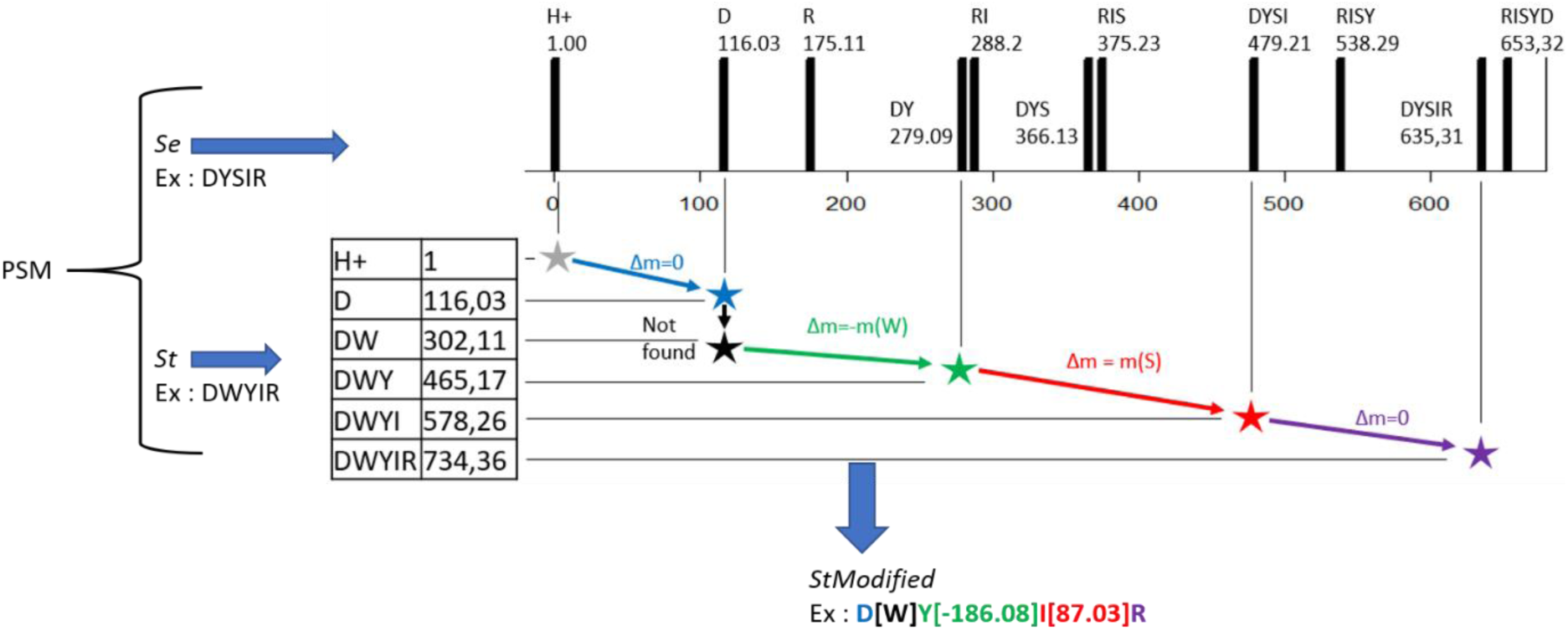
Overview of SpecGlob. As an example, the PSM (DYSIR, DWYIR) is taken as input in SpecGlob. Peptide DYSIR plays the role of the experimental spectrum *Se* while peptide DWYIR plays the role of its best identification, called *St*. The *St* spectrum contains the masses of the *b*-ions after fragmentation of the *St* peptide. As *Se* plays the role of a spectrum obtained from an unknown peptide, it contains both the masses of the *b*-ions and *y*-ions after fragmentation of the peptide (displayed in black vertical lines). To perform the best alignment of masses (according to a user defined score system), SpecGlob fills a dynamic programming table according to rules described in Equation (1) (Materials and Methods). The backtrack step returns the *StModified* string, which can then be used to retrieve the original *Se* peptide. In our example, *StModified* is interpreted as follows: D is aligned to *Se* without any mass offset, W is not found in *Se*, Y is aligned but the alignment requires a mass offset of - 186.08 Da (which is the mass of W), I is aligned with a mass offset of 87.03 Da (which is the mass of S), and R is aligned without any mass offset. Thus, starting from *StModified*, it is possible to retrieve the *Se* sequence DYSIR using a post-processing algorithm which will infer both the deletion of W and the insertion of S after Y.

### 2.4 MODPlus (version v1.02, parameters setting in Supplementary Data, section 3)

MODPlus reports several PSMs per spectrum, ranked by decreasing probability; we selected only the first one to compute statistics.

### 2.5 Execution time measurement

Software have been executed on a laptop equipped with an Intel i7 (2.6 GHz) with 16 Gb of memory allocated to the JVM, running under Windows 10.

### 2.6 Post-processing algorithms (detailed in Supplementary Data, section 4)

Following SpecGlob, post-processing algorithms allow, based on *StModified*, to (1) classify the modifications according to their degree of interpretability (green/orange/red modifications); (2) classify the PSMs in a similar fashion; and (3) transform green modifications into amino acid sequences.

## 3. Results

SpecGlob takes as input a list of PSMs, and considers them one after the other. Note that, at this point, such list of PSMs – with one or several PSMs per spectrum – can be provided as input for SpecGlob by any OMS method.

For each PSM, the best alignment found by SpecGlob is provided as a string, that we call *StModified*, which uses a specific syntax: (1) if two consecutive masses corresponding to the mass of an amino acid are aligned with two masses of *Se* without any insertion of mass offset, this amino acid is reported as such in *StModified*; (2) if the mass difference corresponding to an amino acid is found between masses in *Se*, but the alignment of these masses requires a mass offset, then this amino acid is written in *StModified*, followed by the value of this mass offset between brackets; (3) finally, if the mass difference corresponding to one amino acid is not used in the alignment, this amino acid is written between brackets in *StModified*. By design, for each PSM the sum of all the mass offsets inserted in the alignment is equal to Δ*m*. Several alignments are illustrated Table 1.

**Table 1:** Examples of *StModified* strings provided by SpecGlob, and their interpretations.

| *Se* | *St* | *StModified* | #m | From *St* to *Se* | *Se?* |
| --- | --- | --- | --- | --- | --- |
| GVTACCITK | GITACCITK | G[I]T[-14,02]ACCITK | 1 | $m(I) - 14.02 = m(V)$<br>→ Substitute(I,V) | Y |
| EGASDEWIR | EASDEWIR | EA[57,02]SDEWIR | 1 | $57.02 = m(G)$<br>→ Insert(G) | Y |
| DYSIR | DWYIR | D[W]Y[-186,08]I[87,03]R | 2 | $186.08 = m(W)$<br>→ Deletion(W)<br>$87.03 \text{ Da} = m(S)$<br>→ Insert(S) | Y |
| VCASIYQK | VSFVIFVVIPIH<br>ASIYGAK | [V][S][F][V][I][F][V]V<br>[-791,46] [I][P][I][H]A<br>[-300,25]SIY[G][A]K | 3 | $791.46 = m(VSFVIFV)$<br>→ Delete (VSFVIFV)<br>$m(IPIH-300.25) = m(C)$<br>→ Substitute (IPIH, C)<br>$m(GA) = m(Q)$<br>→ Substitute (GA, Q) | Y |
| QVSVIQWSSIVH<br>GEQCCSVWNAK | QVSVIAK | QVSVIA[1957,82]K | 1 | $1957.82 \text{ Da} = ?$ | N |

Each row displays a PSM (*Se*, *St*) provided by SpecOMS, as well as its output *StModified* by SpecGlob and its interpretation. For readability, spectra (*Se* and *St*) are represented by their corresponding peptides: although *Se* plays the role of an unknown spectrum, its peptide sequence can be subsequently used to validate our results. The *StModified* column shows the strings returned by SpecGlob, which provides an optimal alignment of *St* to *Se*. Information contained in *StModified* is transformed by a series of editing operations (deletions, insertions or substitutions) described in the fifth column, whenever this is possible. The #m column displays the number of modifications in *StModified*. The rightmost column indicates whether the correct *Se* sequence can be retrieved after the editing operations.

In order to validate SpecGlob, we interpreted two simulated spectra datasets carrying systematic modifications on two or three specific amino acids by SpecOMS and SpecGlob on the one hand, and by MODPlus on the other hand. Because spectra were simulated, we were able to compute the percentage of spectra for which each approach restores the exact sequence with its induced modifications (Table 2), a *sine qua non* condition to consider *a posteriori* identifications as correct. Our first remark concerns the execution time of the combination SpecOMS/SpecGlob, which is particularly fast compared to MODPlus. When the search space is limited to the most frequent known modifications (a parameter set in MODPlus), SpecOMS/SpecGlob is about 40 times faster, a ratio that increases to 90 when all the modifications from Unimod are allowed.

**Table 2:**
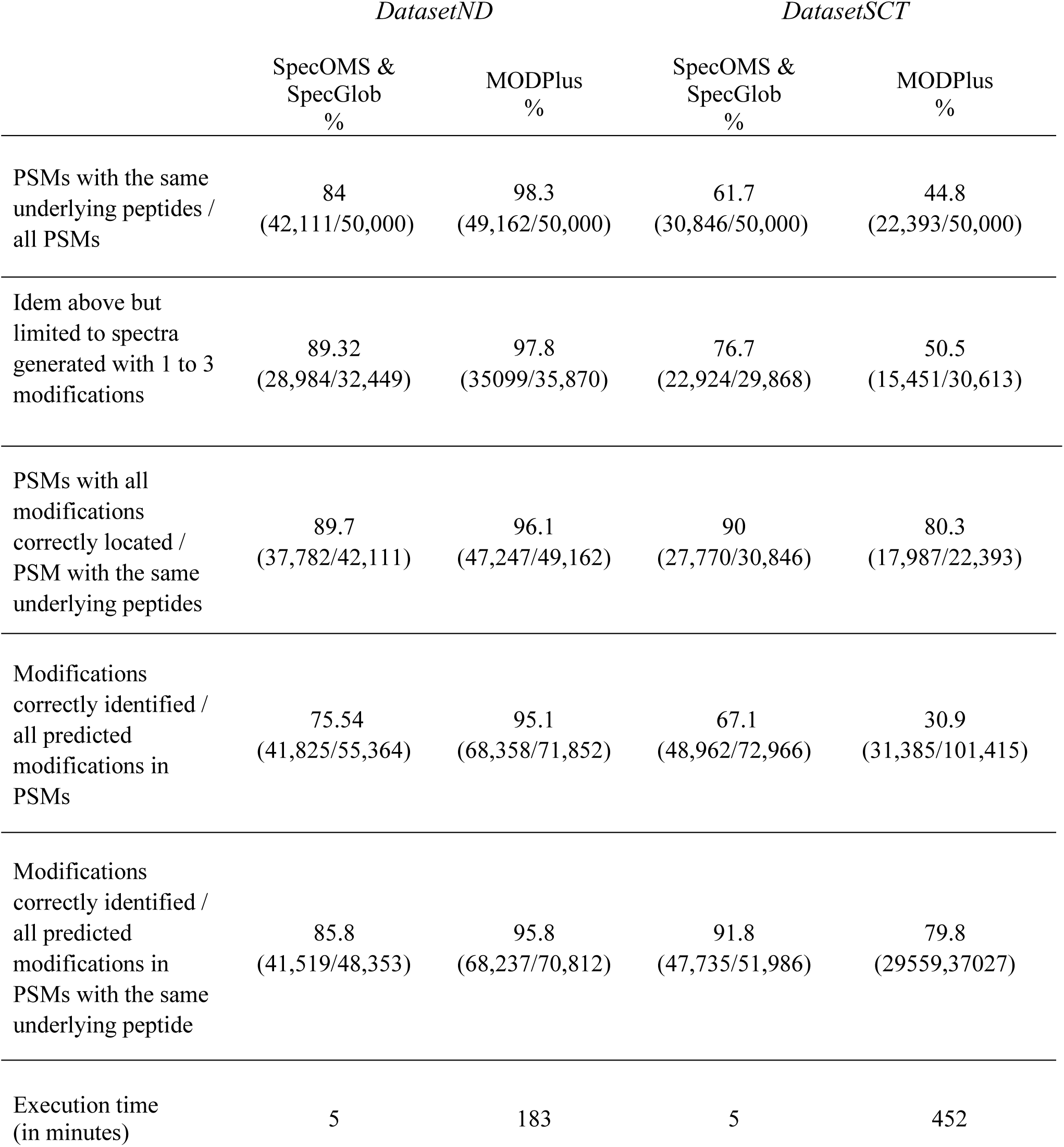
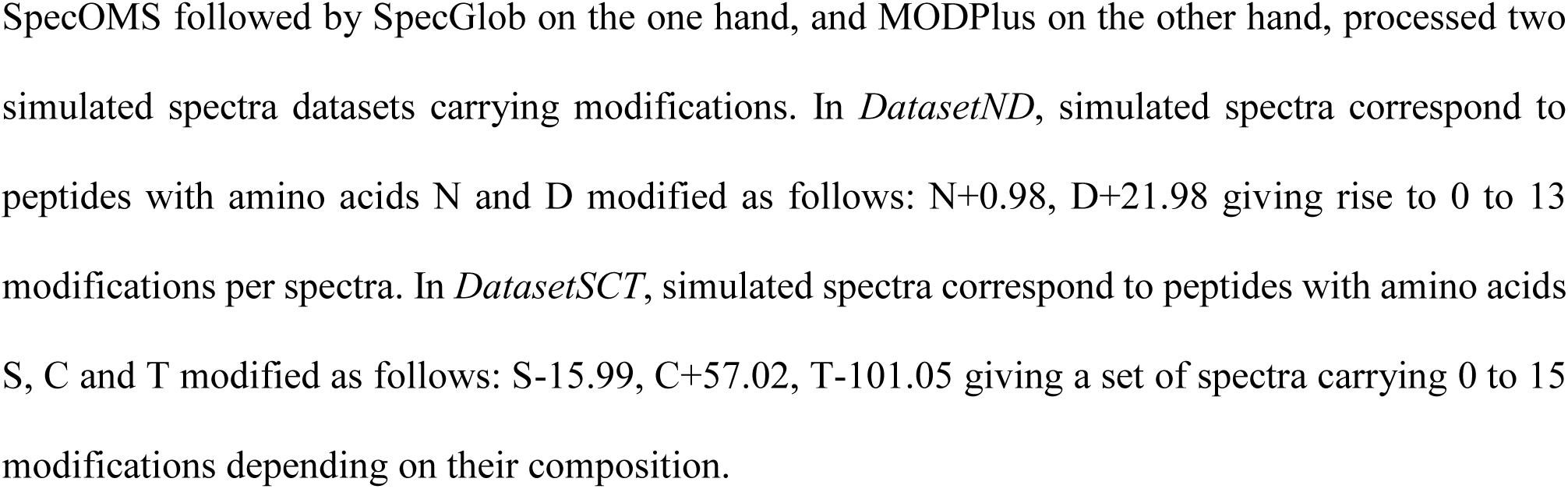
Comparison between SpecOMS/SpecGlob and MODPlus on two simulated spectra datasets.

Concerning the ability of each approach to correctly interpret modified spectra, MODPlus performs extremely well on *DatasetND*, while SpecOMS/SpecGlob is better at discovering unexpected modifications on *DatasetSCT* (Table 2). This result is not surprising. Indeed, *DatasetND* has been designed to be an ideal spectra dataset for MODPlus, which selects candidate peptides that share sequence tags with *Se*, and next performs the search with an input modification list (limited to Unimod or to a user-defined list). In *DatasetND*, only two abundant modifications perfectly described in Unimod were simulated on peptides. By contrast, it should be noted that SpecOMS/SpecGlob processes data without any *a priori*. On the other hand, *DatasetSC*T contains three modifications among which the deletion of T, a modification unreferenced in Unimod. The absence of amino acid deletions in Unimod explains 96% of the incorrect PSMs provided by MODPlus (spectra representing peptides with at least one T), while other misinterpretations are mainly due to the presence of several consecutive modifications close to each other that MODPlus tends to interpret as one.

Hence, although MODPlus is more efficient than SpecOMS/SpecGlob when it comes to analyzing abundant and known modifications in samples (such as *in DatasetND*), SpecOMS/SpecGlob is better at quickly highlighting the large variety of modificationsa sample can carry (including unknown modifications). Moreover, it should also be noted that SpecOMS/SpecGlob obtain approximately the same correct identification rates on both datasets, which shows a certain stability in its behaviour. Secondly, since SpecGlob can quickly align PSMs without any *a priori*, we challenged it on its ability to identify multiple and complex modifications, possibly involving several amino acids. For this, we processed a set of 455,404 PSMs returned by SpecOMS (David *et al*. 2017) when all theoretical spectra generated from the human proteome were pairwise compared, while excluding self-identifications (*AllvsAll* dataset). We believe that only a simulated spectra dataset such as *AllvsAll* allows an evaluation of the accuracy of the results (percentage of well-localized offsets) in such a complex situation.

While about a quarter of the PSMs can be optimally aligned with only one modification – that can involve one or several amino acids-, a large majority of PSMs yields an optimal alignment carrying *strictly more* than one modification. Once SpecGlob has aligned spectra and produced a *StModified* string, a post-processing algorithm can be invoked to interpret the values of offset masses as editing operations. We introduced a colour classification, in the same spirit as in (Lysiak *et al*. 2021), to partition PSMs processed by SpecGlob into three categories, each category reflecting a certain degree of complexity in retrieving the amino acid sequence that generated *Se*, starting from *StModified*. A modification is considered

- *green* if it can be unambiguously converted into an editing operation: insertion of one amino acid, substitution of one amino acid by another or deletion of one or several consecutive amino acid(s);
- *orange* if it can be explained as an editing operation, but where some ambiguity remains: in the case of an insertion of several amino acids whose identities are known, but for which their order in the sequence remains uncertain. Note that because we need to limit execution time, we limited each insertion or substitution to a sequence of at most *three* amino acids;
- *red* when a modification is neither green nor orange.

We extended our colour classification from the *modification* level to the *PSM* level. A *green* PSM contains only *green* modifications; an *orange* PSM contains only *green* or *orange* modifications (and at least one is *orange*); a PSM is *red* otherwise. The number of PSMs in each category, summarized Table 3, establishes that SpecGlob is capable of interpreting a large proportion of PSMs returned by SpecOMS, although several editing operations differentiate *Se* from *St*. Moreover, even if a PSM is not green (and cannot be completely interpreted), we found that the length of the longest amino acid subsequence that can be unambiguously interpreted is increased on average by two amino acids when the post-processing step is applied on *StModified*, thus increasing the information extracted from the PSMs in the orange or red categories.

**Table 3:** Colour distribution of modifications and PSMs on Dataset *AllvsAll*.

|  | Green | Orange | Red | Total |
| --- | --- | --- | --- | --- |
|  | 740,458 | 286,616 | 390,639 |  |
| #Modif. | 232,925 in green<br>PSMs | 141,132 in orange<br>PSMs | 390,639 in red<br>PSMs | 1,417,713 |
|  | 152,332 in orange<br>PSMs | 145,484 in red<br>PSMs |  |  |
|  | 355,201 in red<br>PSMs |  |  |  |
| #PSMs | 132,137 | 114,454 | 208,813 | 455,404 |

## 4. Discussion

Because considering the alignments of all experimental spectra to all theoretical spectra remains unrealistic due to excessive computational load, OMS methods have developed different strategies to filter and align PSMs. MODPlus is representative of those methods that only consider known modifications during alignments. If this strategy is well adapted to the analysis of experimentally enriched samples, the number of correct interpretations drops as soon as the number of spectra carrying unknown modifications increases. It is an important drawback since it is known that reference modification databases are partial (preparation of samples can infinitely vary and generate unexpected modifications) and are, and will probably remain, unsuitable for a certain number of studies (studies of modifications induced by food processes, for example).

Other methods, such as SpecOMS or MSFragger (Kong *et al*. 2017), filter the most promising PSMs (pairs of spectra that already share a certain number of peaks) and next, interpret the resulting Δ*m* between *Se* and *St* as a single modification, which may lead to an incorrect suggestion of modification. Once a PSM references the correct peptide, SpecGlob is efficient at finding the best alignment (around 92% of modifications correctly located on *DatasetSCT*). The percentage of correct PSMs, which seems to be the weak point, could still certainly be improved. Firstly, note that one spectrum in *DatasetSCT* may carry many modifications (up to fifteen in this dataset). The number of mass offsets required for an optimal alignment is not defined or limited in advance in SpecGlob, but when this number is too high, performances rapidly decline. For instance, selecting only spectra that carry less than four modifications is a good way to improve confidence (see Table 2). Secondly, we set SpecOMS so as to provide only one PSM per *Se* as input to SpecGlob. Delivering more than one PSM per *Se* and then selecting the ‘best’ *St* returned by SpecGlob (considering e.g. the score of the obtained alignment and/or the colour of the corresponding PSM) would also certainly increase the number of correct PSMs. Given the rapidity of SpecGlob, increasing the number of processed PSMs is perfectly conceivable.

We also demonstrate, when processing *AllvsAll* dataset, that even in the case where the *Se* sequence is not completely restored, SpecGlob followed by its post-processing step, can partly restore it. More precisely, SpecGlob can produce subsequences of the amino acids that are present in *Se*, thus giving clues to peptide identification – and in turn to protein identification – with an efficiency that depends on their length and composition.

To put this in perspective, we believe that, whatever strategy is adopted to generate the PSMs, SpecGlob is useful to confirm or suggest well evidenced modifications by aligning spectra very quickly without preconceived ideas. The way we write *StModified* is useful in itself to summarize alignments, because the information it contains goes beyond location of the modifications, by highlighting stretches of amino acids effectively detected in the spectrum. An alternative is to graphically visualise the spectra, together with annotated masses, which should give access to more details for the user, but is highly time consuming. Besides, *StModified* strings are easy to handle by various scripts, which allows to gather information for further investigations, as we did for instance when applying our colour classification.

## 5. Conclusions

Our study shows that SpecGlob is a promising tool for interpretation of mass spectra through post-analysis of PSMs. SpecGlob aligns pairs of spectra from collections of PSMs returned by OMS methods, possibly introducing several mass offsets, in order to increase the quality of the initial alignments. By running a fast competition between several possible alignments of spectra within a PSM, SpecGlob is likely to propose alternative interpretations supported by well-aligned spectra to those that might have been anticipated by users. The results we obtained on the interpretation of theoretical spectra should now be used for testing and using SpecGlob on experimental spectra, possibly adapting it for that purpose. Its expected relevance on experimental spectra is supported by its very good behaviour, which we demonstrated in this paper. Besides, SpecGlob supplements OMS methods by providing interpretation of spectra under the form of a handy alignment that is easy to interpret.

## Abbreviations

OMS: Open Modification Search
PSM: Peptide Spectrum Match
SPC: Shared Peaks Count

## Author contributions

Albane Lysiak: conceptualization (equal), Methodology (equal), software, formal analysis, writing, review and editing (equal); Guillaume Fertin: conceptualization (equal), Methodology (equal), review and editing (equal); Géraldine Jean: conceptualization (equal), Methodology (equal), review and editing (equal); Dominique Tessier: conceptualization (equal), Methodology (equal), writing, review and editing (equal).

## Conflict of Interest declaration

The authors declare that they have no affiliations with or involvement in any organization or entity with any financial interest in the subject matter or materials discussed in this manuscript.

## Funding

Supported by the French National Research Agency (ANR-18-CE45-004), ANR DeepProt.

## Data access statement

The human proteome was downloaded from Ensembl 99, release GrCh38 on the Ensembl FTP server. Proteins predicted with the annotation “protein coding” were added to the contaminant proteins downloaded from the cRAP contaminant database http://ftp.thegpm.org/fasta/cRAP/. The SpecOMS software is available at https://github.com/dominique-tessier/SpecOMS.

## Notes

### Competing Interest Statement

The authors have declared no competing interest.

